# Host contact dynamics shapes richness and dominance of pathogen strains

**DOI:** 10.1101/428185

**Authors:** Francesco Pinotti, Éric Fleury, Didier Guillemot, Pierre-Yves Böelle, Chiara Poletto

## Abstract

The interaction among multiple microbial strains affects the spread of infectious diseases and the efficacy of interventions. Genomic tools have made it increasingly easy to observe pathogenic strains diversity, but the best interpretation of such diversity has remained difficult because of relationships with host and environmental factors. Here, we focus on host-to-host contact behavior and study how it changes populations of pathogens in a minimal model of multi-strain interaction. We simulated a population of identical strains competing by mutual exclusion and spreading on a dynamical network of hosts according to a stochastic susceptible-infectious-susceptible model. We computed ecological indicators of diversity and dominance in strain populations for a collection of networks illustrating various properties found in real-world examples. Heterogeneities in the number of contacts among hosts were found to reduce diversity and increase dominance by making the repartition of strains among infected hosts more uneven, while strong community structure among hosts increased strain diversity. We found that the introduction of strains associated with hosts entering and leaving the system led to the highest pathogenic richness at intermediate turnover levels. These results were finally illustrated using the spread of *Staphylococcus aureus* in a long-term health-care facility where close proximity interactions and strain carriage were collected simultaneously. We found that network structural and temporal properties could account for a large part of the variability observed in strain diversity. These results show how stochasticity and network structure affect the population ecology of pathogens and warns against interpreting observations as unambiguous evidence of epidemiological differences between strains.

**Author summary:** Pathogens are structured in multiple strains that interact and co-circulate on the same host population. This ecological diversity affects, in many cases, the spread dynamics and the efficacy of vaccination and antibiotic treatment. Thus understanding its biological and host-behavioral drivers is crucial for outbreak assessment and for explaining trends of new-strain emergence. We used stochastic modeling and network theory to quantify the role of host contact behavior on strain richness and dominance. We systematically compared multi-strain spread on different network models displaying properties observed in real-world contact patterns. We then analyzed the real-case example of *Staphylococcus aureus* spread in a hospital, leveraging on a combined dataset of carriage and close proximity interactions. We found that contact dynamics has a profound impact on a strain population. Contact heterogeneity, for instance, reduces strain diversity by reducing the number of circulating strains and leading few strains to dominate over the others. These results have important implications in disease ecology and in the epidemiological interpretation of biological data.

## Introduction

Interactions between strains of the same pathogen play a central role in how they spread in host populations. [1–7]. In *Streptococcus pneumoniae* and *Staphylococcus aureus*, for instance, several dozen strains can be characterized for which differences in transmissibility, virulence and duration of colonization have been reported in some cases [8, 9]. Strain diversity may also affect the efficacy of prophylactic control measures such as vaccination or treatment. Indeed, strains may be associated with different antibiotic resistance profiles [3, 5, 10, 11], and developed vaccines may only target a subset of strains [2, 3, 12]. With the increasing availability of genotypic information, it has become easy to describe the ecology of population of pathogens and to monitor patterns of extinction and dominance of pathogen variants [13–17]. However, the reasons for multi-strain coexistence patterns (e.g. coexistence between resistant and sensitive strains) or dominance of certain strains (e.g. in response to the selection pressure induced by treatment and preventive measures) remain elusive. One may invoke selection due to different pathogen characteristics, but also environmental and host population characteristics, leading to differences in host behavior, settings and spatial structure may affect the ecology of strains [14–19]. In particular, human-to-human contacts play a central role in infectious disease transmission [20]. This is increasingly well described thanks to extensive high-resolution data - including mobility patterns [21–23], sexual encounters [24], close proximity interactions in schools [25, 26], workplaces [27], hospitals [16, 28–31], etc.-, that enable basing epidemiological assessment on contact data with real-life complexity [32, 33]. For instance, the frequency of contacts can be highly heterogeneous leading more active individuals to be at once more vulnerable to infections and acting as super-spreaders after infection [24, 33–35]. Organizational structure of certain settings (school classes, hospital wards, etc.) and other spatial proximity constraints lead to the formation of communities that can delay epidemic spread [36, 37]. Individual turnover in the host population is also described as a key factor in controlling an epidemic [20, 38]. It is likely that, since they impact the spread of single pathogens, the same characteristics could affect the dynamics in multi-strain populations. It was shown, indeed, that network structure impacts transmission with two interacting strains [39–46], the evolution of epidemiological traits [47–49] and the effect of cross-immunity [50, 51]. Yet in these cases, complex biological mechanisms - such as mutation, variations in transmissibility and infectious period, cross immunity - were used to differentiate between pathogens, thereby making the role of network characteristics difficult to assess in its own right.

For this reason, we focused on the dynamical pattern of human contacts and examined whether it contributes to shaping the population ecology of interacting strains under minimal epidemiological assumptions regarding transmission. We described a neutral situation where all strains have the same epidemiological traits and compete via mutual exclusion (concurrent infection with multiple strains is assumed to be impossible) in a Susceptible-Infected-Susceptible (SIS) framework. We studied the spread of pathogens in a host population during a limited time window, disregarding long-term evolution dynamics of pathogens. More precisely, new strains were introduced through host turnover rather than *de novo* mutation or recombination in pathogens. We quantified the effect of network properties on the ecological diversity in strain populations with richness and dominance indicators. We assessed in turn heterogeneities in contact frequency, community structure and host turnover by comparing simulation results obtained with network models exhibiting a specific feature. We then interpreted *S. aureus* carriage in patients of a long-term care facility in the light of these results.

## Results

### Effects of contact heterogeneity

We simulated the stochastic spread of multiple strains on a dynamical contact network of individuals (nodes of the network). Individuals can be either susceptible or infected with a single strain at a given time, and, for each strain, *β* and *µ* indicate the transmission and the recovery probability respectively. We assumed continuous turnover of individuals, who enter the system with probability *λ*_in_, and associated injection of previously unseen strains, carried by incoming individuals with probability *p_s_*. In order to probe the effect of contact heterogeneity on strain ecology we compared a homogeneous model (HOM) in which all nodes have the same activity potential, i.e. they have equal rate of activation to establish contacts, with a heterogeneous model (HET) where the average activity potential is the same, but the rate of activation is heterogeneous across individuals [34]. Then, for each network model we characterized the structure of pathogen population at the equilibrium through ecological diversity measures, including species richness and evenness/dominance indices [52, 53].

We show sample epidemic trajectories in Fig 1A and average quantities in panels B-D of Fig 1. The prevalence, summarized in Fig 1B for different transmissibility values, displays a well-known behavior for both static and dynamic networks: contact heterogeneities lower the transmissibility threshold above which total prevalence is significantly above zero, thus allowing the spread of pathogens with low-transmissibility. At the same time, however, heterogeneities hamper the epidemic spread when *β* is large, reducing the equilibrium prevalence [35]. Fig 1 shows that richness (i.e. the number of distinct strains co-circulating) is not linked to the prevalence in a straightforward way. For sufficiently large 1, the reduction in richness of HET with respect to HOM is important even for the case with mild contact heterogeneity, when prevalence is barely affected (Fig 1C). The scaling between prevalence and richness is not linear as *β* varies (Fig 1D), and the relation between the two quantities varies appreciably among contact networks. In correspondence of a fixed value of prevalence, heterogeneous networks have lower richness - e.g. a prevalence value of 250 corresponds to 21% lower richness in HET with respect to HOM, as highlighted in Fig 1D. This fact can be explained by the balance between injection of new strains and extinction of already circulating ones. The extinction of a stochastic SIS process is certain, being the disease-free state the unique absorbing state. When multiple SIS processes spread on the same network, the persistence time of a single process is short, in the sense that it scales linearly with the size of the system [54] (in contrast to the lifetime of a single SIS process which scales exponentially with the size of the system when *β* is above the threshold value [55]). Here we find that network heterogeneity shortens the persistence time of a strain (see also Fig S1 in the supporting information). Indeed active nodes involved in a larger number of contacts get infected more frequently [35]. Strains introduced by low-activity nodes are likely to be surrounded by nodes already infected, thus limiting transmission. As a consequence they encounter extinction more easily. In other words, contact heterogeneities strengthen the competition induced by mutual exclusion.

**Fig 1.**
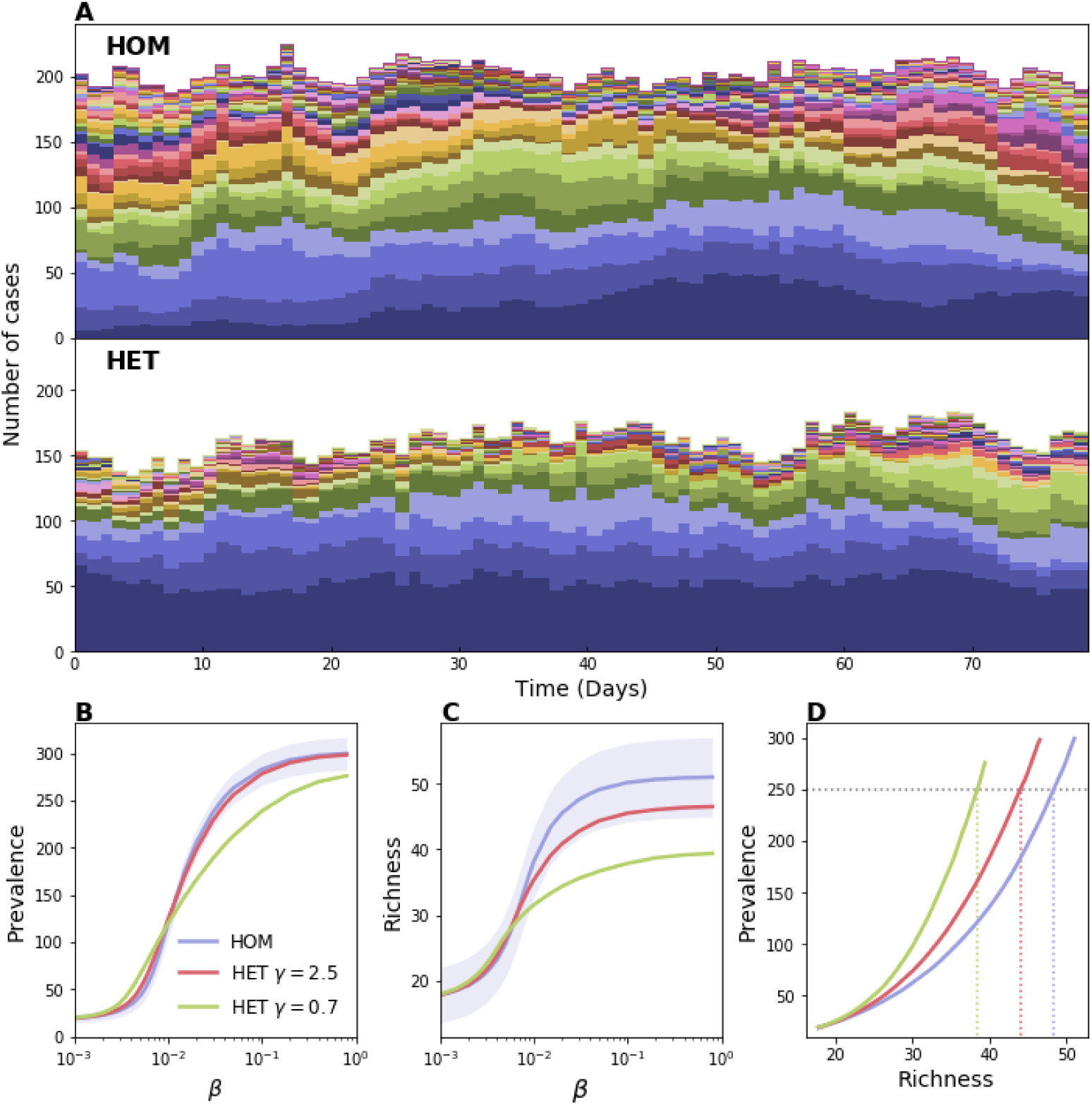
Effect of contact heterogeneity on strain richness. Comparison between a homogeneous (HOM) and a heterogeneous (HET) network. In HOM each node activates with probability *a*_H_ = 0.285 and the network average degree is 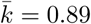. In HET the activation probability of each node is drawn randomly from a power-law distribution with the same average activity of HOM. *γ* is the exponent of the power-law, thus lower values of *γ* correspond to a higher contact heterogeneity. We chose a population of 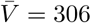 individuals, an average length of stay *τ* = 10 days, the probability an incoming individual brings a new strain *p_s_* = 0.079 and *µ* = 0.00192 (see Material and methods). (A) Sample time series of strain abundance for HOM and HET with *γ* = 0.7, here *β* = 0.02. (B) Average prevalence vs *β*. Two levels of heterogeneity are here considered for HET. For the sake of visualization, the shaded area corresponding to the standard deviation is shown only for HOM. (C) Average richness vs *β*. (D) Average prevalence vs richness. Dashed lines are shown as a guide to the eye, highlighting variation in richness induced by network topology.

The presence of hubs not only reduces richness for sufficiently large *β*, but affects more profoundly the distribution of strains’ abundances, i.e. the strain-specific prevalence, leading to stronger fluctuations (Fig 2A). If on one hand hubs accelerate the extinction of certain strains, on the other hand they act as super-spreaders and amplify the prevalence of other strains. This results in a situation of dominance where certain strains, despite having no biological advantage, become able to overcome the others reaching a significant proportion of the population. This situation is synthesized by the Berger-Parker index, defined as the relative abundance of the most abundant strain - an alternative indicator, the Shannon evenness, is shown in Fig S2 in the Supplementary Information. Fig 2B-C shows how this quantity varies when increasing strain transmissibility. As expected, at low *β* values the short transmission chains produced by different strains barely interact. The competition becomes, instead, more pronounced as *β* increases and, consistently, the effect of the network topology becomes more relevant.

**Fig 2.**
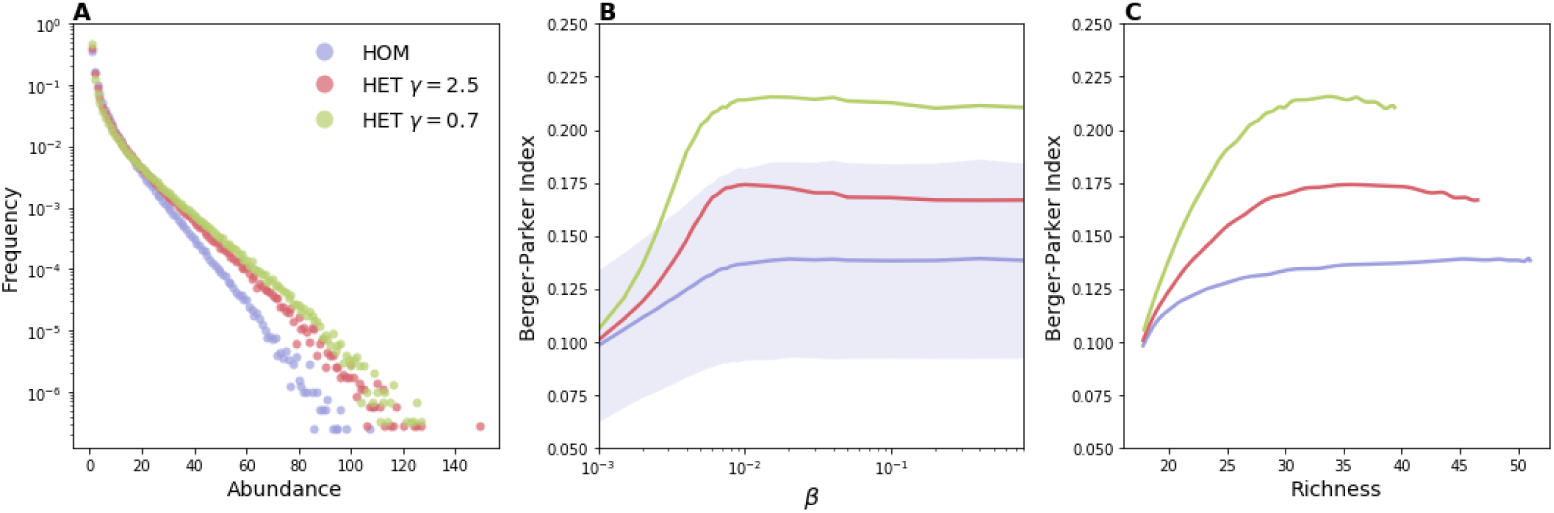
Effect of contact heterogeneity on strain dominance. (A) Abundance distribution. (B) Berger-Parker index vs *β*. Berger-Parker index is defined as the relative abundance of the most abundant strain. For the sake of visualization, the shaded area corresponding to the standard deviation is shown only for HOM. (C) Berger-Parker index vs richness. Parameters are the same as in Fig 1.

We tested whether additional mechanisms of strain injection were leading to different results. In Fig S3 we assumed new strains to infect susceptible nodes already present in the system with probability *q_s_*, mimicking in this way transmissions originating from an external source, as it can happen in real cases. The plot of Fig S3 shows the same qualitative behavior described here.

### Effect of community structure

We considered a community model (COM) with *n_C_* communities in which all nodes are as active as in HOM, but direct a fraction *p_IN_* of their links within their community and the rest to nodes in the remaining *n_C_* − 1 communities. The closer *p_IN_* is to 1, the stronger the repartition in communities.

Fig 3A,B shows that a network with communities displays a higher richness for large *β*; even when community structure barely affects prevalence (Fig 3B). However, the effect is important only when communities are fairly isolated (*p_IN_* = 0.99) and the injection from the outside is not so frequent - otherwise the effect is masked by strain injection which occurs uniformly across communities. In particular, for the values of *p_IN_* = 0.78 and *p_s_* = 0.079, chosen to match the real-case scenario discussed latter in the text (i.e. the spread *S. aureus* within a hospital), the difference with the homogeneous case is very small. For low *β*, the behavior of the Berger-Parker index follows the trend in richness. The initial decrease in this indicator is due to the increase in richness, that occurs at constant prevalence and is thus associated to a decrease in the average abundance [56] - green curve in Fig 3C corresponding to *p_IN_* = 0.99 and *p_s_* = 0.01. At larger values of *β*, instead, increased competition levels induced higher dominance levels.

**Fig 3.**
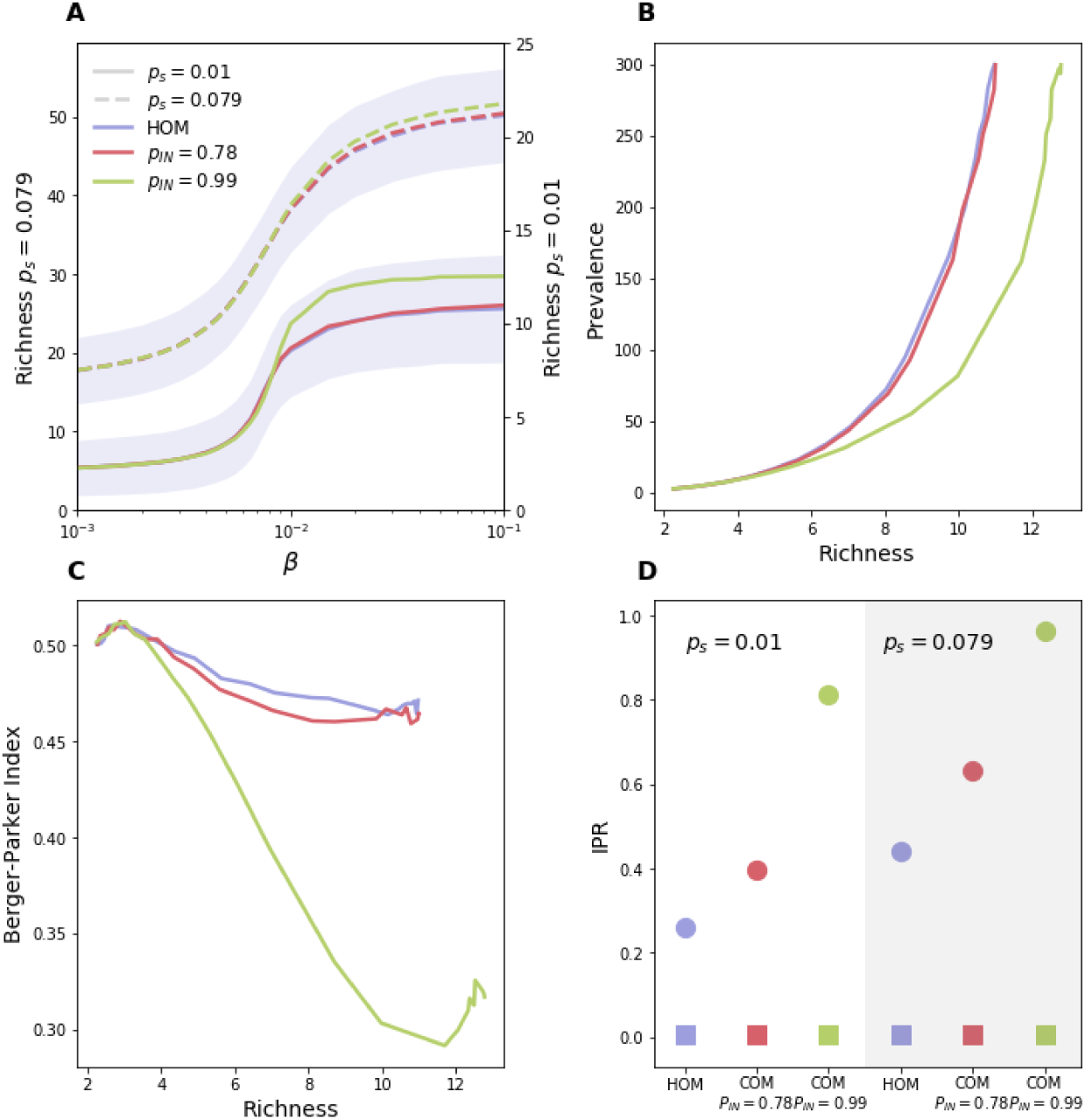
Impact of community structure. (A) Richness vs *β* for HOM (blue lines) and for COM with *p_IN_* = 0.78 (red lines) and *p_IN_* = 0.99 (green lines). For both COM models we have set *n*_C_ = 6. Solid lines correspond to *p_s_* = 0.01, while dashed lines correspond to *p_s_* = 0.079. (B) Prevalence vs richness for *p_s_* = 0.01. (C) Berger-Parker index vs richness for *p_s_* = 0.01. (D) Average IPR for both *p_s_* = 0.01 (white background) and *p_s_* = 0.079 (gray background) and for *β* = 0.02. Squares correspond to IPR obtained from total prevalence while circles correspond to IPR obtained from strains’ abundances. Here no injection due to transmission from an external source is assumed (*q_s_* = 0). The effect of this second mechanism is shown in Fig S3.

The increase in strain diversity is due to the reduced competition among strains introduced in different communities. When coupling among communities is low, indeed, strains may spend the majority of time within the community they were injected in, thus avoiding strains injected in other communities. Fig 3D confirms this hypothesis by showing the Inverse Participation Ratio (*IPR*) [57] that quantifies uniformity in the repartition of abundance across communities. Values close to zero indicate uniform repartition, while, conversely, values close to 1 indicate that, on average, a strain is confined within a single community for most of the time (more details are reported in the Material and Methods section). The strength of the community structure does not affect the repartition of the total prevalence (squares in the plot), however it alters the average *IPR* value computed from the abundance of single strains, thus strains become more localized as *p_IN_* increases. Notice that a certain degree of localization is present also in the homogeneous network, due to the case in which injected strains cause very few generations before getting extinct.

### Effect of turnover of individuals

Another important factor is node turnover as it has a profound impact on the ecological dynamics of strains for two reasons: incoming individuals contribute to richness by injecting new strains; on the other hand, the removal from the population of infected nodes breaks transmission chains and hampers the persistence of strains. The result of the interplay between these two mechanisms is summarized by the plot of richness as a function of *β* and node length of stay, *τ*, - Fig 4A. The figure, obtained with the HOM model, shows two distinct regimes. In the former case, richness decreases as *τ* increases, because replacement of individuals becomes slower and injections less frequent. In the high *β* regime, instead, the average richness at fixed *β* does not depend monotonically on the node turnover but it is instead maximized at intermediate *τ*. Interestingly, the optimal value of *τ* decreases as *β* increases. This behavior can be explained by looking at the balance between injection and extinction that determines the equilibrium value of richness, 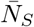. This reads [58]:

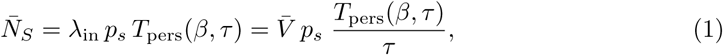

where *λ*_in_ *p_s_* is the rate at which new strains are introduced and *T*_per_*_s_* is the average persistence time of a strain. The trade-off between injection and extinction appears as the ratio between the two time scales, *T*_per_*_s_* and *τ*. In the limit *τ* → 0 the spread plays no role, even for high *β*. As *τ* increases, newly introduced infectious seeds have a higher probability to spread, thus the average extinction time initially increases super-linearly with *τ* (see Fig S4 in the Supplementary Information) resulting in an increase of richness. However, past a certain value of *τ*, *T*_per_*_s_* does not grow super-linearly anymore, thus a further increase in *τ* is detrimental for pathogen diversity because it is associated to fewer introductions. This general behavior was not altered by the accounting for introductions by transmissions from an external source as shown in Fig S3.

**Fig 4.**
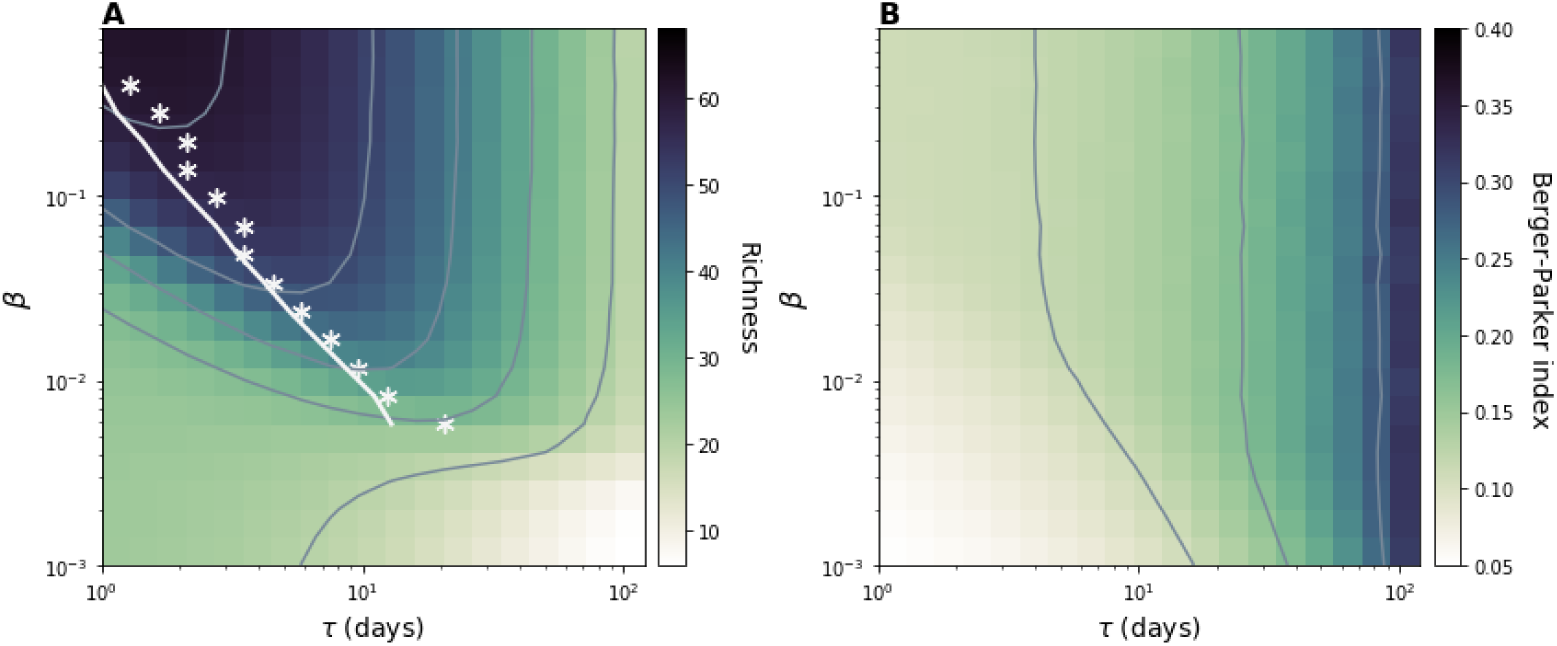
Effects of node length of stay on strain diversity. (A) Average richness and (B) Average Berger-Parker index. Contour plots are shown in both figures. While exploring *τ* we also set the value of the average network size 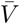 to 306. For each value of *β* we highlight in panel (A) the value of the length of stay corresponding to the maximum richness (white asterisks) together with the analytical prediction (white line). Here *µ*= 0.00192, 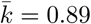, *a*_H_ = 0.28.

We derive an approximate formula for *T*_pers_ considering an emerging strain competing with a single effective strain formed by all other strains grouped together. This formulation, enabled by the neutral hypothesis, makes it possible to write the master equation describing the dynamics and using the Fokker-Planck approximation to derive persistence times (see Material and Methods). Analytical results well reproduce the behavior recovered by simulations, and, in particular, the value of the length of stay maximizing richness for different *β* as shown by the comparison between white stars and continuous line in Fig 4C. The quantitative match for other values of *p_s_* is reported in Fig S5 in the Supplementary Information.

Unlike richness, Berger-Parker index always increases monotonically with the length of stay - Fig 4B. This behavior is due to the correlation of this indicator with average abundance, similarly to what we discussed in the previous section.

### Spread of *S. aureus* in a hospital setting

We conclude by analyzing the real-case example of the *S. aureus* spread in a hospital setting [10, 59]. We used close-proximity-interaction (CPI) data recorded in a long-term health-care facility during 4 months by the i-Bird study [16, 28, 31]. These describe a high-resolution dynamical network, whose complex structure reflects the hospital organization, the subdivision in wards and the admission and discharge of patients [60]. Together with the measurements of contacts, weekly nasal swabs were done to monitor the *S. aureus* carriage status of the participants and identify the spa-type and the antibiotic resistance profile of the colonizing strains.

The modeling framework considered here well applies to this case. The SIS model is widely adopted for modeling the *S. aureus* colonization [62, 63], and the assumption of mutual exclusion is made by the majority of works to model the high level of cross-protection recognized by both epidemiological and microbiological studies [64, 65]. The dynamic CPI network was previously shown to be associated with paths of strain propagation [16]. Consistently, we assumed that transmission is mediated by network links with transmissibility *β*. In addition, new strains are introduced in the population carried by incoming patients, or contacts with persons not taking part in the study.

Fig 5A shows weekly carriage and its breakdown in different strains. Prevalence and richness fluctuate around the average values 87, 3 ± 6, 3 cases and 39, 8 ± 2 strains, respectively. Simulation results are reported in Fig 5B, that displays the impact of transmission and introduction rate on richness and prevalence. When introduction rate is low we find a positive trend between richness and prevalence, consistently with the synthetic case. For higher injection rates, instead, the relation between richness and prevalence becomes first less pronounced and then decreasing because as soon as the transmission rate becomes larger the injection is hampered by the depletion of susceptibles.

**Fig 5.**
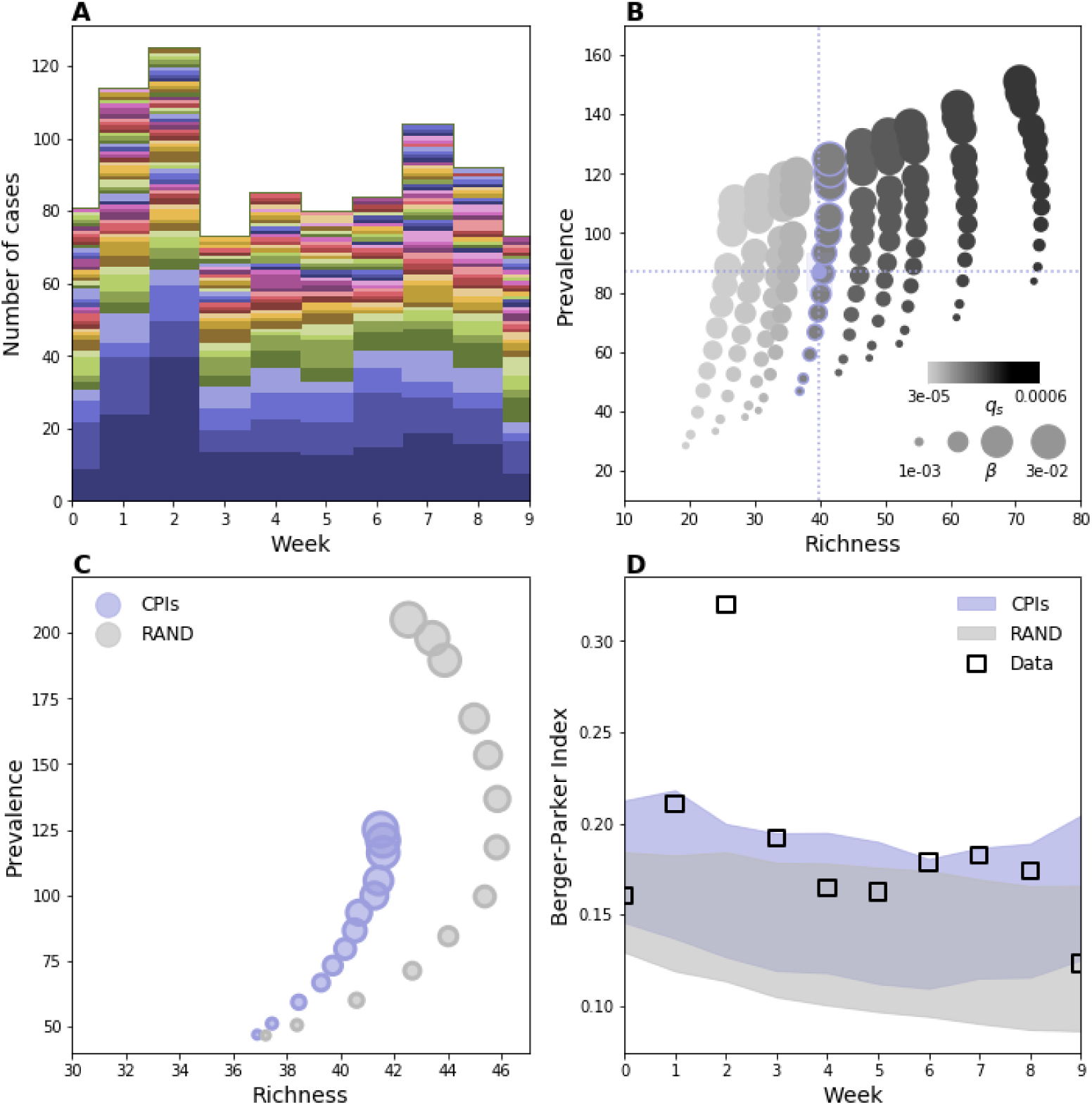
*S. aureus* population structure on hospital network. (A) Weekly carriage data. (B) Prevalence vs. richness from simulations on the hospital network for different *β* and different rate of introduction, here tuned by the parameter *q_s_*. Blue dashed lines represent the average empirical values. (C) Prevalence vs richness for hospital contact data (blue dots) and RAND (grey dots). The hospital curve corresponds to the curve in (B) with blue-contour markers. (D) For each week, Berger-Parker index in carriage data along with the same quantity from the simulations. The shaded areas indicate the average plus/minus the standard deviation obtained from 1000 stochastic runs. For each network, parameter values are the ones that reproduce empirical prevalence and richness. Duration of colonization is assumed here to be 35 days [61]. Alternative values of this parameter led to same qualitative results (Fig S6).

To quantify the effect of contact patterns on *S. aureus* population ecology we compared simulation results with the ones on a network null model. Specifically we built the RAND null model that randomizes contacts while preserving just the first and the last contact of every individual. The randomization preserves node turnover, number of active nodes and links and destroys contact heterogeneities and community structure along with higher-order correlations. Fig 5C shows the comparison for different transmissibility values. The effect of the network is consistent with the theoretical results described for a heterogeneous network, i.e. smaller richness values correspond to the same prevalence in the real network compared to the homogeneous one. We then quantified the level of dominance of the multi-strain distribution by means of the Berger-Parker index. We chose for each network introduction and transmissibility rates that better reproduce empirical richness and prevalence and, interestingly, we found that, for the two cases, same average richness and prevalence correspond to different Berger-Parker behaviors. The Berger-Parker obtained with the real network is the highest and the one that better matches the empirical values - i.e. the empirical values are within one standard deviation of the mean for almost all weeks. Based on this result we argue that contact heterogeneities, along with the other properties of the contact network, contribute to the increased dominance of certain strains.

## Discussion

Multiple biological and environmental factors concur in shaping pathogen diversity. We focused here on the host contact network and we used a minimal transmission model to assess the impact of this ingredient on strain population ecology, quantifying the effects of three main network properties, i.e. heterogeneous activity potential, presence of communities and turnover of individuals. Results show that the structure and dynamics of contacts can alter profoundly strains’ co-circulation. Contact heterogeneities, by quickly driving low-abundance strains to extinction, reduce strain richness and favor strain dominance. Highly active nodes are known to play an important role in outbreak dynamics by acting as super-spreaders [33]. Here we showed that a similar mechanism could allow strains with no biological advantage to generate a large number of cases and outcompete other equally fit strains. This mechanism may potentially bias the interpretation of biological data. Dynamical models that do not properly account for contact structure could overestimate the difference in strains’ epidemiological traits in the attempt to explain observed fluctuations in strain abundance induced in reality by super-spreading events. Moreover, these models could provide biased assessment of transmission vs. introduction rates.

The presence of communities causes the separation of strains and mitigates the effect of competition thus enhancing co-existence. A similar behavior was already pointed out for the spread of *S. pneumoniae*, as induced by age assortativity [66], for the case of S. aureus where distinct settings were considered [62], and for a population of antigenic distinct strains in presence of cross-immunity [51]. We found that the impact of community structure is not so strong, and it is likely minor when individuals of different communities have frequent contacts. No appreciable variation was observed, indeed, for *p*_IN_ = 0.78, chosen to match the inter-ward coupling of the hospital CPIs network. Similar results can be expected for school classes or workplace departments presenting a similar level of community mixing. The effect on richness becomes appreciable for low community coupling (e.g. *p*_IN_ = 0.99 in Fig 3). This is consistent with a certain degree of diversity observed among strain belonging to separated communities, as it is the case of different hospitals [15].

Eventually, the analysis of turnover of individuals revealed major effects on strain diversity, when this mechanism is also the main responsible for the introduction of strains in the population. When transmissibility is low richness decreases with host length of stay. When transmissibility is above the epidemic threshold we showed the existence of an optimal value of the length of stay that maximizes strain richness as a result of the interplay between two competing timescales, namely the typical inter-introduction time and the average persistence time of a strain. This provides insights for the spread of bacterial infections in transmission settings, such as hospitals or farms, that are of particular relevance for the spread of antimicrobial resistance and that are characterized by a rapid host turnover [15, 31, 67]. For the case of hospitals, for instance, they suggest that variations in patients’ length of stay, as induced by a change of policy, could have appreciable effects on the population structure of nosocomial pathogens.

We adopted a neutral model to better disentangle the relative role of the different network properties. A wide disease-ecology literature addressed the consequences of neutral hypotheses on multi-strain balance in order to provide a benchmark for interpreting the observed co-existence patterns and gauging the effect of selective forces potentially at play [11, 18, 68, 69]. Many of these works addressed, for instance, the co-existence between susceptible and resistant strains of *S. pneumoniae* [11, 68]. However, this assumption was rarely adopted in network models, that consider for the majority strains with different epidemiological traits with the aim of describing pathogen selection and evolution [47–49, 70]. Strains were assumed to have the same infection parameters in [50, 51], where the role of community structure and clustering was analyzed in conjunction with cross-immunity. With respect to these works, the minimal transmission model used here enabled a transparent comprehension of the role of the network. Multiple identical SIS processes can be mapped, in fact, on a single SIS process, in such a way that the wide literature of single SIS processes allows for a better understanding of the behavior recovered in the simulations [32, 33]. Strains can be also grouped in two macro strains. This strategy allowed us to adopt the viewpoint of an emerging strain and study its competition with the others seen as a unique macro-strain. The associated Markov equation and Fokker-Planck approximation allow computing the average extinction time, capturing the key aspects of the dynamics. We focused here on three major properties of human contacts. Future work can leverage on a similar transmission model to address other properties known to alter spreading dynamics, such as heterogeneous inter-contact time distribution or topological and temporal correlations.

As a case study, we analyzed the spread of *S. aureus* in a hospital taking advantage of the simultaneous availability of contact and carriage information [16]. The temporal and topological features of the network lead to a lower prevalence and richness with respect to the homogeneous mixing (although the effect was quite small). In addition, similar prevalence and richness values are associated to different dominance levels in different networks - i.e. different values of the Berger-Parker index -, with the real network leading to a higher dominance as observed in reality. This behavior can be explained by the theoretical results and can be attributed essentially to the effect of contact heterogeneities, considering that the community structure does not have appreciable effects for this network, as discussed above. The importance of accounting for host contacts and hospital organization in the assessment of bacterial spread and designing intervention has been recognized by several studies [16, 28–31, 61]. Here we show that this element may be critical also for understanding the population ecology of the bacterium. It is important to note however that, while the realistic network provides results that are closer to the data, this ingredient explains only part of the heterogeneity observed in the abundance. This shows that the contact network is a relevant factor, but other factors should be considered as well. The approach used here is intentionally simplified, as we focused on the main dynamical consequences of the contact network. Clearly, more detailed models can be designed to reproduce more closely the data. A certain degree of variation in the epidemiological traits could be at play, as for example the fitness cost of resistance [8]. Role of hosts in the network (e.g. patients vs. health-care workers), and heterogeneities in health conditions, antibiotic treatment and hygiene practices are also known to affect duration of carriage and chance of transmission [16, 28, 31, 61]. Eventually, we must consider that the comparison of model output with carriage data is also affected by the limitation of the dataset itself, already described in [16]. In particular, the weekly swabs may leave transient colonization undetected. Moreover, while the relevance of CPIs as proxies for epidemiological links has been demonstrated [16], the transmission through the environment (e.g. in the form of fomites) is also possible.

The understanding provided here can be relevant for other population settings, temporal scales and geographical levels. In addition, the modeling framework could be applied to pathogens other than *S. aureus*, such as *human papillomavirus, S. pneumoniae* and *Neisseria meningitidis*, for which the strong interest in the study of the strain ecology is justified by the public health need for understanding and anticipating trends in antibiotic resistance, or the long-term effect of vaccination [1, 2, 4, 5]. With this respect, if the simple framework introduced here increases our theoretical comprehension of the multi-strain dynamics, more tailored models may become necessary according to the specific case. In particular, we have considered complete mutual exclusion as the only mechanism for competition. In reality, a secondary inoculation in a host that is already a carrier may give raise to alternative outcomes, such as co-infection or replacement [71]. In addition infection or carriage may confer a certain level of long-lasting strain-specific protection and/or a short-duration transcendent immunity [11, 50]. Eventually mechanisms of mutation and/or recombination are at play and their inclusion into the model can be important according to the time scale of interest.

## Materials and methods

### Network models

For all the network models considered, the stochastic generative algorithms return a sequence of time-stamped networks and share a similar general scheme:

**Turnover dynamics:** new nodes arrive according to a Poisson process with rate *λ*_in_ and leave after a random time drawn from an exponential probability distribution with average *τ*. After a short initial transient, population size is Poisson distributed with average 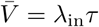.

**Activation Pattern:** during each time step each node activates with a given probability that depends on the actual model considered. Each active node then creates a number of stubs which is drawn from a zero-truncated Poisson distribution. The active status lasts for just a single time step.

**Stub matching:** stubs are then matched according to the actual model considered.

The generative models considered in this work are:

**HOM:** in this model each node has the same probability *a*_H_ to be active during each time step. Stubs are matched completely at random in order to form links. We discard eventual self-links and multiple-links that may occur during the matching procedure.

**HET:** here each node has its own activation probability *a_i_*, drawn from a power-law distribution *P* (*a*) ∝ *a*^−γ^, with *a* ∈ (*ϵ*, 1]. We tune the variance varying *γ* - lower *γ* higher variance. We then set *ϵ* to have the average activity *ā* equal to *a*_H_ in HOM. Stub-matching procedure is the same as in HOM.

**COM:** incoming nodes are assigned to one among *n_C_* communities with equal probability, in such a way that communities have the same size on average. Each stub is matched within the respective community with probability *p_IN_* or outside the community with complementary probability.

### Hospital network and null models

We use a dynamical contact network obtained from CPI data collected during the i-Bird study in a French hospital. Details of the network are already reported in [16]. We aggregated the CPIs daily keeping the information about their cumulated duration within each day. We discard CPIs relative to the first 2 weeks and the last 4 weeks of dataset, corresponding to a period of adjustments in the measurements and progressive dismissal of the experiment, respectively. Simulations conducted with the CPIs network were compared with results obtained with a null model which we refer to as RAND. According to this randomization scheme the activity of node is randomized while respecting the constraint that removal and addition of contacts must not alter the time of the first and the last contact of each node (*t_S_* and *t_L_* respectively). Notice that RAND preserves the number of nodes that are present at any time in the network by preserving their first contact *t_S_* and their length of stay *t*_L_ – *t_S_*. Null models randomizing the latter properties lead to misleading results when node length of stay is heterogeneous and node turnover occurs [72]. RAND also sets all contact weights equal to the average weight value.

### Simulation details

Transmission dynamics is entirely stochastic and emerges from the combination of transmission, recovery and strain injection. During time step *t* each node is updated according to its state: each infected node transmits the strain it is carrying to a susceptible neighbor with probability *β*, and infected nodes turn susceptible with probability *µ*. For the *S. aureus* case study, transmission probability depends the cumulated duration of the contact within the day (*w_ij_*) according to the expression 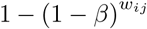. Due to mutual exclusion, an individual can be infected by a single strain at a given time [73]. New unseen strains are injected with rate *ι*. At each time step incoming individuals can bring a new strain with probability *p_s_*. In addition, susceptible individuals may turn infectious carrying a new strain with probability *q_s_*. The two mechanisms mimic respectively incoming infectious individuals (e.g. admission of colonized patients) and transmission from an external source (in the hospital example this corresponds to contacts with individuals that were not participating in the study). Injection rate is thus given by 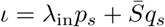, where 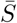 is the number of susceptible at the equilibrium. In the theoretical analysis in the main paper we assumed *q_s_* = 0 for simplicity, thus variations in *ι* where induced by variations in *λ*_in_ and *p_s_*. The case *q_s_ >* 0 was considered in the Supplementary Information. In the hospital case study *p_s_* was set to 0.079 (directly estimated from the data), while *q_s_* was explored.

In the theoretical analysis parameters were set, in the majority of cases, to match the hospital case study - e.g value of *p_s_*, average number of nodes 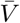, average activation rate *ā*, number of communities (*n_C_*), etc.

### Analysis of carriage data

Carriage data was obtained from weekly swabs in multiple body areas, including the nares. Swabs that resulted positive to *S. aureus* were further examined. Spa-type and antibiotic resistance profiles (MSSA or MRSA) were then determined. In this work we regard two strains as different if they differ in spa-type and/or antibiotic resistance profile. We considered carriage data obtained from nasal swabs dismissing other body areas since the anterior nares represent the most important niche for *S. aureus* [74].

### Ecological measures and other indicators

We described strain population diversity through standard ecological indicators. The abundance of a strain *i*, *N_i_*, is the strain-associated prevalence. From this quantity we computed the abundance distribution, being the frequency of strains with abundance *N*. The Berger-Parker index is the relative abundance of the dominant strain, i.e. 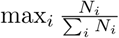.

To analyze repartition of strains across communities we use the Inverse Participation Ration (*IPR*) [57]. Given a vector 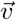 with elements 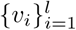, all within [0, 1], the IPR is given by:

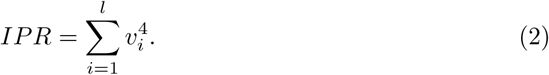

If all the components are of the order (*l*^−1^) then the *IPR* is small. Instead if one component *v_i_* ~ 1 then *IPR ~* 1 too, reflecting localization of 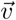. The IPR for total prevalence is computed by setting *v_i_* equal to the fraction of infected individuals belonging to community *i*, while the IPR for a single strain is computed by setting *v_i_* equal to the fraction of individuals infected by that particular strain and belonging to community *i*. We can extend the IPR computation to HOM case by assigning nodes to different groups as in COM but without affecting the stub-matching scheme.

### Analytical results for the homogenous network

In order to estimate the value of the length of stay maximizing the average richness for a given value of *β* when the contact structure is given by the HOM network we consider a homogeneous mixing version of our system.

Due to Eq (1) the calculation of the average richness reduces to the calculation of the average persistence time. In order to estimate such quantity we focus on a particular strain, labelled as “strain A”, which is injected at *t* = 0 and we group all other strains under the label “strain B”. We are allowed to do so because all strains have identical parameters. We therefore reduce our initial, multi-strain problem, to a two-strain problem. Since all new strains that will be injected after *t* = 0 will be labeled as strain B, it is clear that A is doomed to extinction since there exists an infinite reservoir of B. The average time to extinction is therefore the average time to extinction of strain A.

Since HOM network realizes quite well homogeneous mixing conditions we regard our system as homogeneously mixed. Within this framework it is sufficient to specify the numbers of hosts infected by strain A (*m*), hosts infected by strain B (*n*) and susceptible hosts (*s*). The master equation for the joint probability distribution *P* (*m, n, s*) is given by [75]:

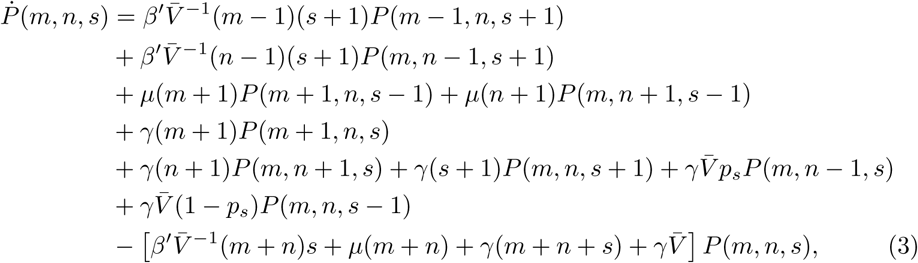

Where 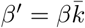. The various terms represent contributions due to infection, recovery, admission and discharge of nodes. In order to obtain some approximate solution to this equation we assume that the average number of individuals *m* + *n* + *s* and the total prevalence *m* + *n* do not fluctuate in time and are therefore equal to 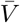 and 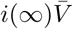 respectively, where *i*(∞) is given by:

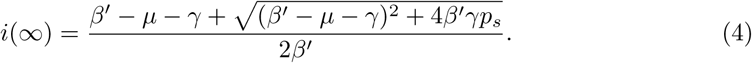

After performing the Van-Kampen size expansion we are left with a Fokker-Planck equation for the density of A 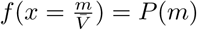:

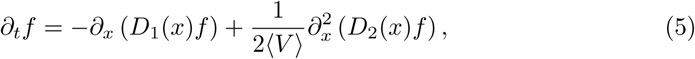

where *D*_1_ = *β*^ʹ^ (1 − *i*(∞)) *x* − *µ* − *γ* and *D*_2_ = *β*^ʹ^ (1 − *i*(∞)) *x* + *µ* + *γ* are the so-called drift and diffusion coefficients respectively.

According to the theory of stochastic processes [75] the average extinction time *T*_per_*_s_*(*x*_0_)(where *x*_0_ represents the initial density of strain A) satisfies:

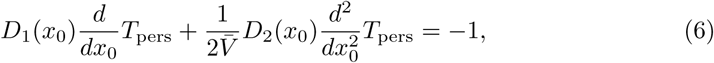

with boundary conditions *T*_per_*_s_*(0)=0 and 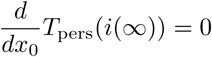. The solution is finally given by:

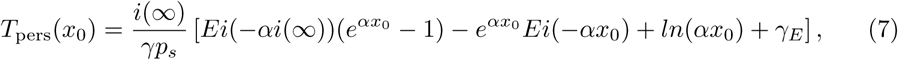

where *Ei*(*x*) is the exponential integral function and *γ_E_* is Euler-Mascheroni constant. When a new strain is introduced its prevalence is just 1, therefore we estimate the average extinction time using 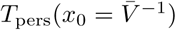.

## Acknowledgments

Authors would like to thank Vittoria Colizza, Lulla Opatowski and Laura Temime for useful discussion.

## Supporting Information Host contact dynamics shapes richness and dominance of pathogen strains

**Fig S1:**
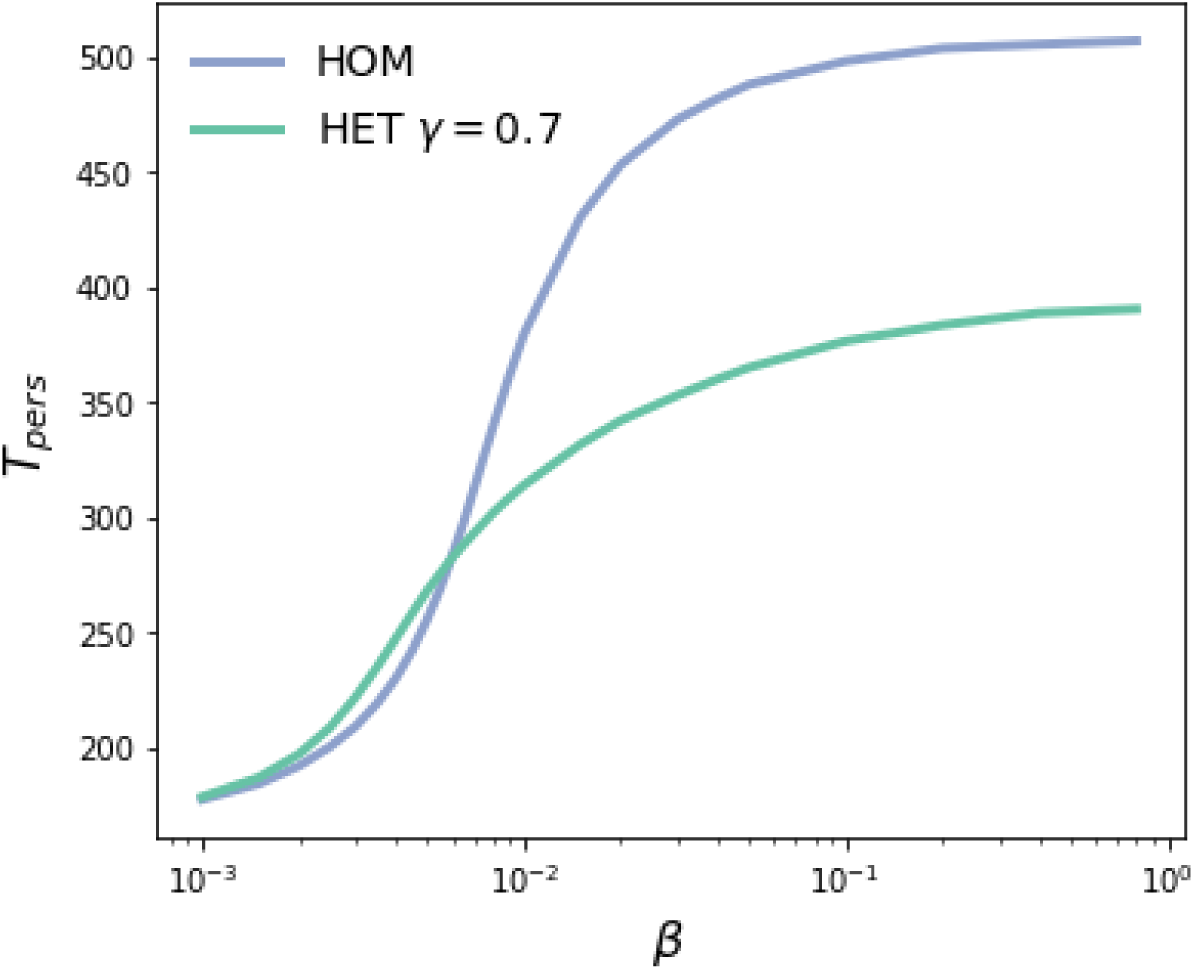
Average extinction times for HOM (blue) and HET with activity distribution exponent *γ* = 0.7 (green) as a function of the transmissibility *β*. We start measuring persistence times once the system has reached a dynamical equilibrium. Parameters are as in Fig 1 in the main text.

**Fig S2:**
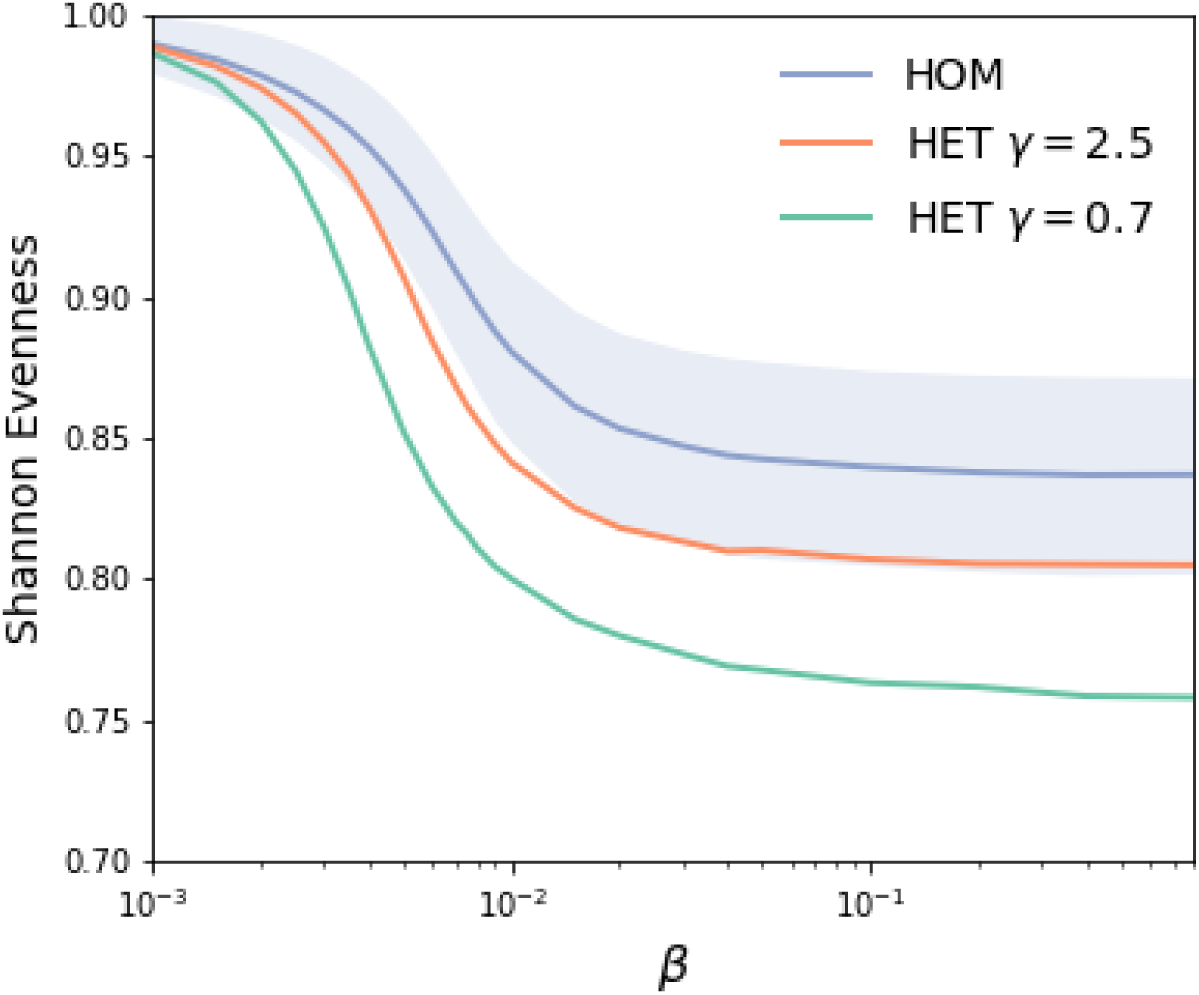
Shannon Evenness for HOM model (blue) and two instances of HET model for activity distribution exponent *γ* = 2.5 (orange) and *γ* = 0.7 (green). Shaded blue area represents standard deviation for HOM. We introduce the relative abundance of the *i*-th strain: 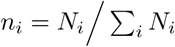, with *N_i_* the abundance of the strain *i* (i.e. the number of infected with strain *i*). Shannon evenness is defined as the normalized Shannon entropy 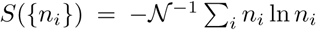, with 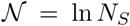. Parameters are the same as in Fig 1 in the main text.

**Fig S3:**
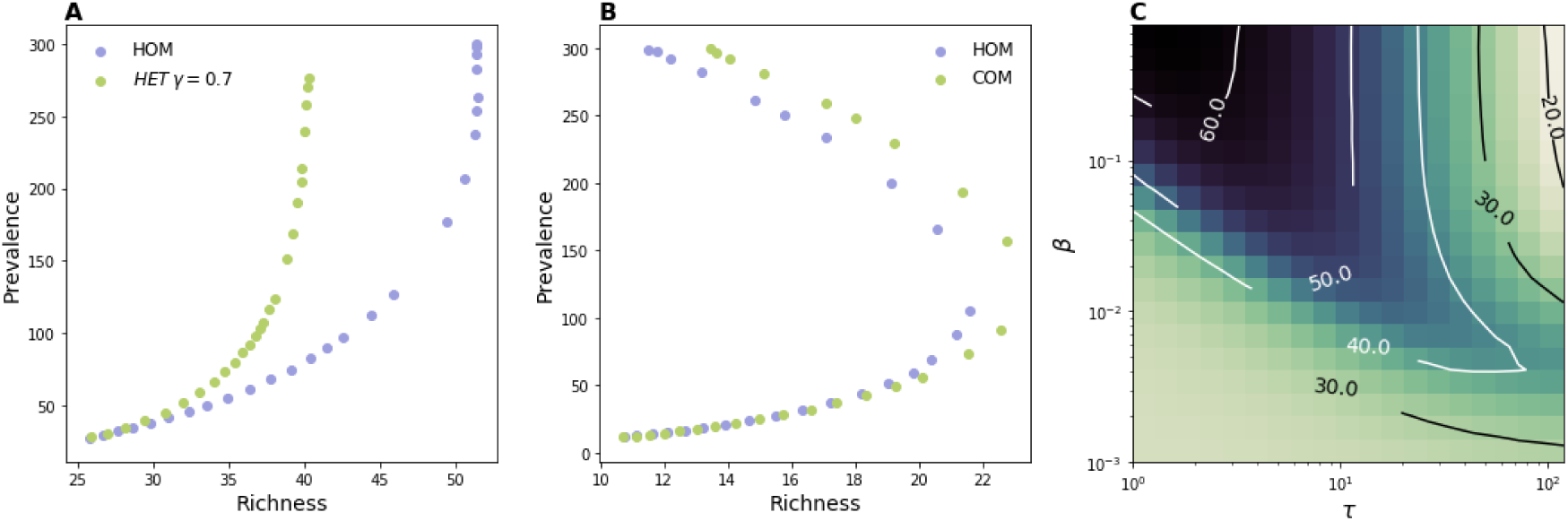
Simulations on various generative network models with transmission from an external source tuned by *q_s_*; here *q_s_* = 0.0002.(A) Richness for HOM model (blue markers) and HET model with activity distribution exponent *γ* = 0.7 (green markers). Here *p_s_* = 0.079. (B) Richness index for HOM model (blue markers) and COM model with within-community connection probability *p_IN_* = 0.99 (green markers). Here *p_s_* = 0.01. (C) Richness as a function of *β* and *τ* for HOM model. Here *p_s_* = 0.079.

**Fig S4:**
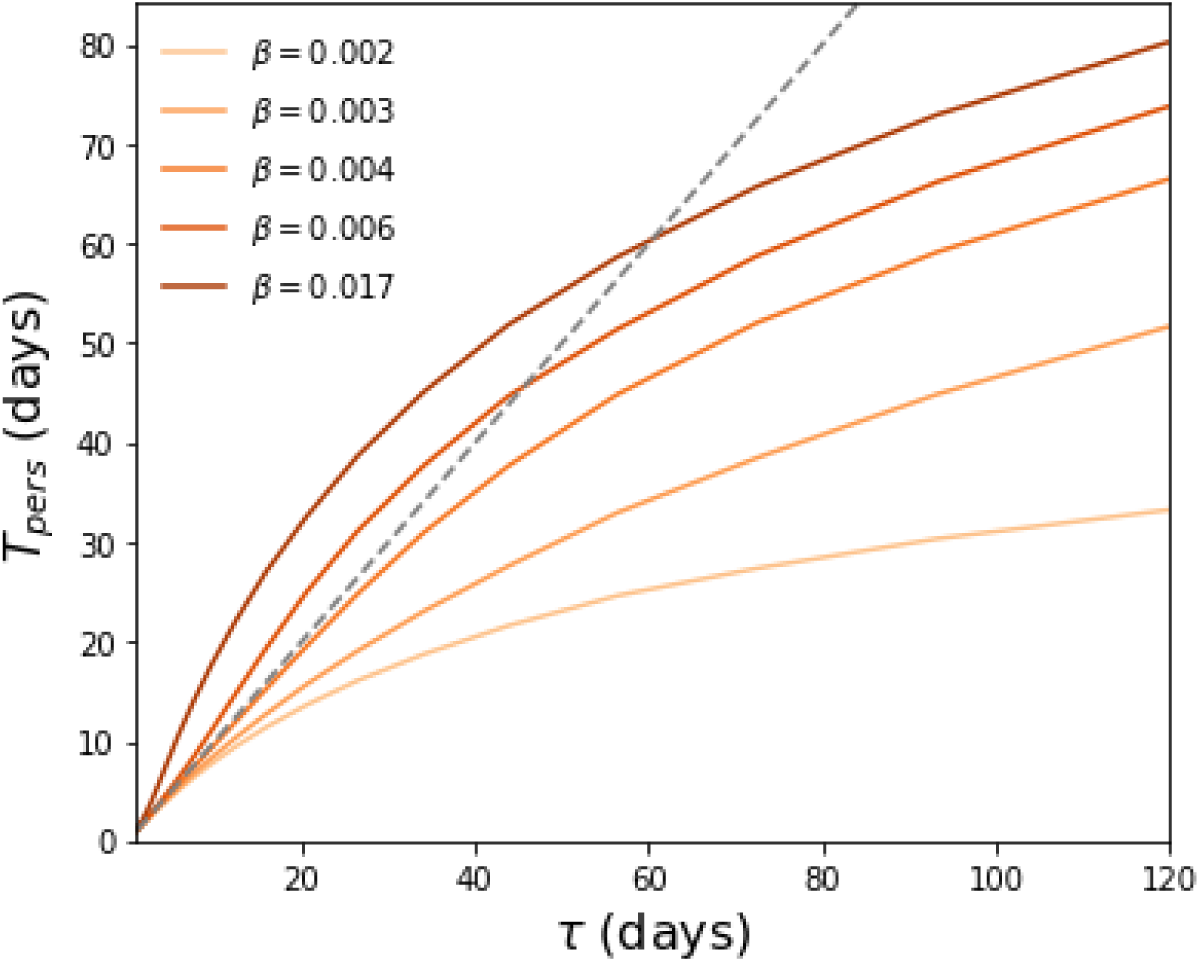
Average persistence time, *T*_pers_, obtained by simulations on HOM model for several values of transmissibility and length of stay. The dashed gray line represents a linear trend as a guide to the eye. Parameters are the same as in Fig 4 in the main text.

**Fig S5:**
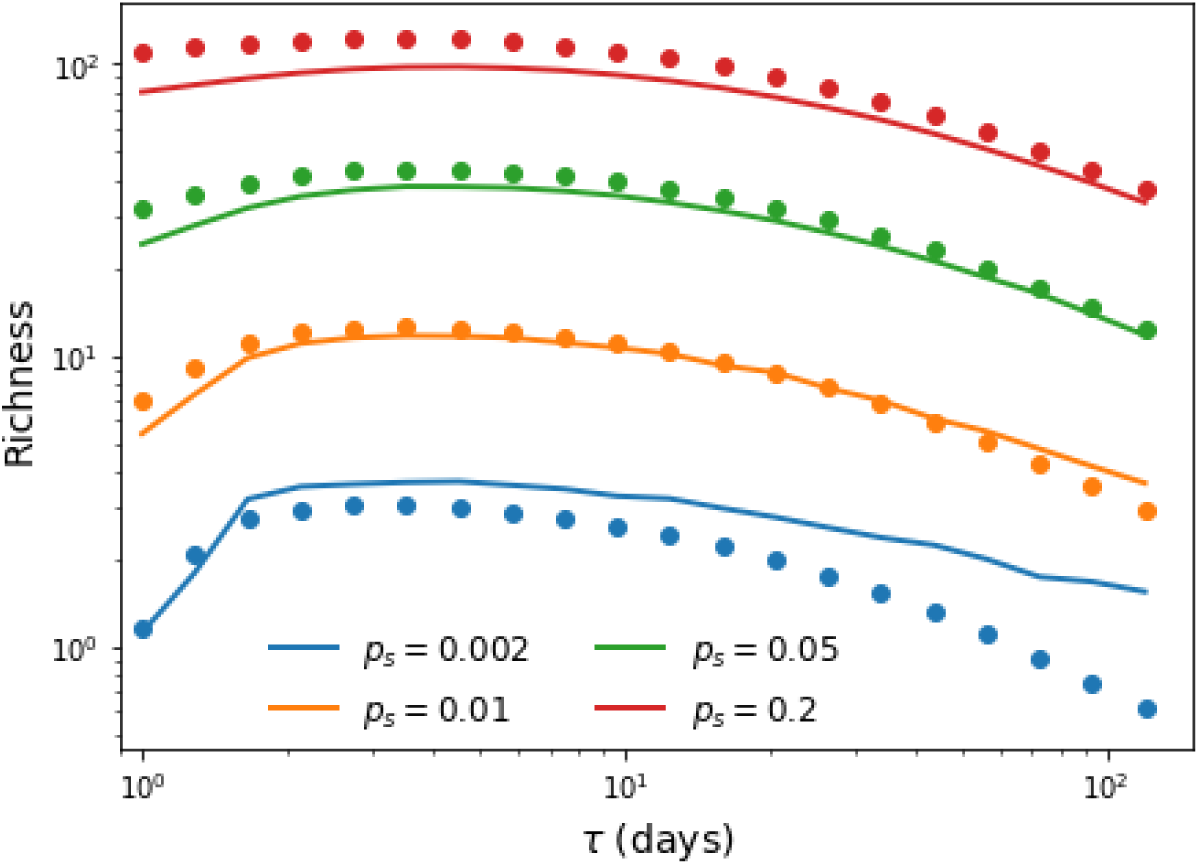
Comparison between simulations for HOM model and analytical predictions obtained using the Fokker-Planck framework introduced in the Materials and Methods section. Solid lines represent average richness obtained by using Eq (1) and Eq (7) from the main text while dots represent simulations results. Here *β* = 0.04 while other parameters are the same as in Fig 4 in the main text.

**Fig S6:**
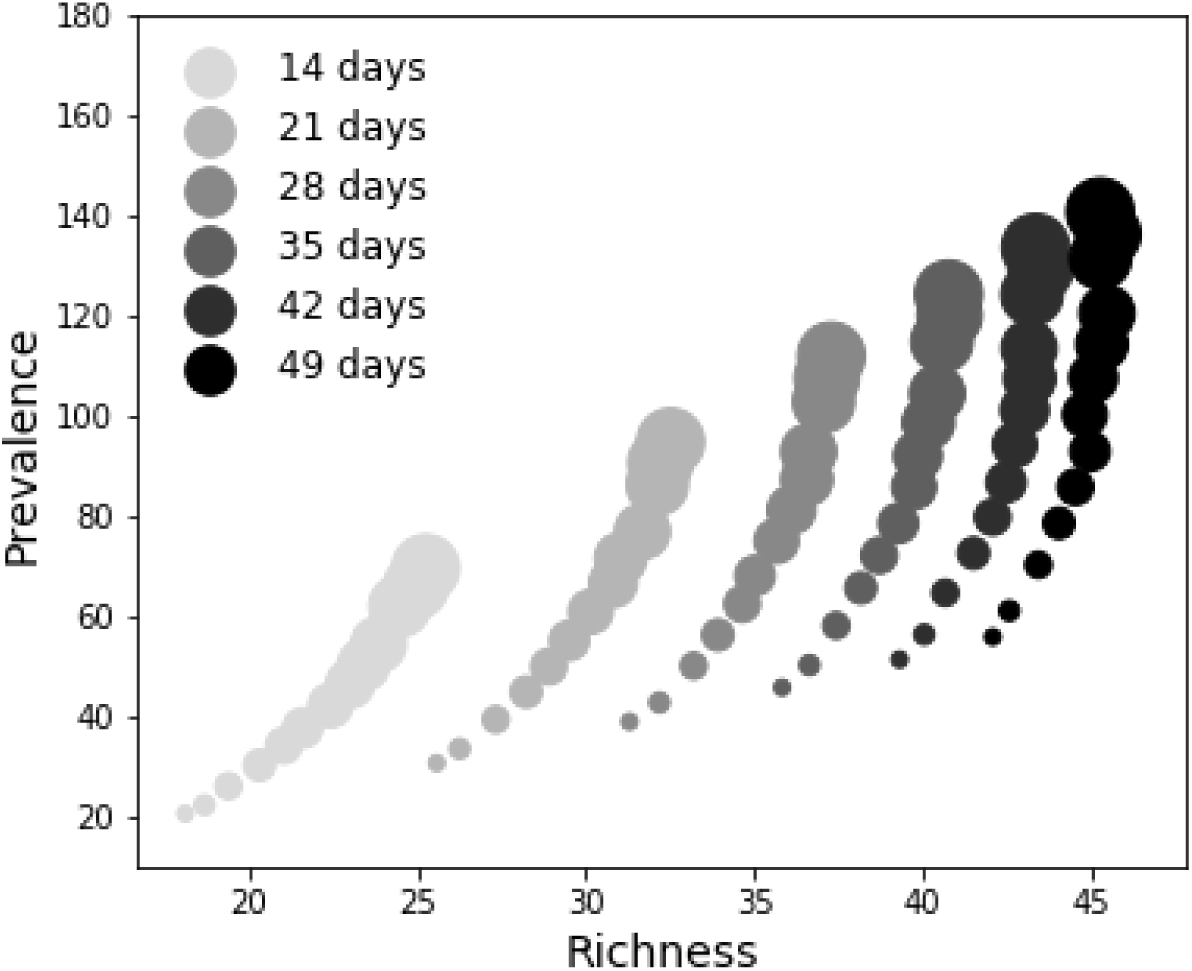
Prevalence vs Richness for several values of the infectious period and using the CPI network. The value of *q_s_* is the same for the curve highlighted in Fig 5B in the main text, *q_s_* = 0.00018. Here dot size is proportional to the magnitude of *β*.

## References

1. Schiffman M, Castle PE. Human Papillomavirus: Epidemiology and Public Health. Archives of Pathology & Laboratory Medicine. 2003;127(8):930–934. doi:10.1043/1543-2165(2003)127¡930:HPEAPH¿2.0.CO;2.

2. Weinberger DM, Malley R, Lipsitch M. Serotype replacement in disease after pneumococcal vaccination. The Lancet. 2011;378(9807):1962–1973. doi:10.1016/S0140-6736(10)62225-8.

3. Bogaert D, de Groot R, Hermans P. Streptococcus pneumoniae colonisation: the key to pneumococcal disease. The Lancet Infectious Diseases. 2004;4(3):144–154. doi:10.1016/S1473-3099(04)00938-7.

4. Atkins KE, Lafferty EI, Deeny SR, Davies NG, Robotham JV, Jit M. Use of mathematical modelling to assess the impact of vaccines on antibiotic resistance. The Lancet Infectious Diseases. 2017;doi:10.1016/S1473-3099(17)30478-4.

5. Chambers HF, Deleo FR. Waves of resistance: Staphylococcus aureus in the antibiotic era. Nat Rev Microbiol. 2009;7(9):629–641. doi:10.1038/nrmicro2200.

6. Reich NG, Shrestha S, King AA, Rohani P, Lessler J, Kalayanarooj S, et al. Interactions between serotypes of dengue highlight epidemiological impact of cross-immunity. J R Soc Interface. 2013;10(86). doi:10.1098/rsif.2013.0414.

7. Cohen T, Colijn C, Murray M. Modeling the effects of strain diversity and mechanisms of strain competition on the potential performance of new tuberculosis vaccines. Proc Natl Acad Sci U S A. 2008;105(42):16302–16307. doi:10.1073/pnas.0808746105.

8. Melnyk AH, Wong A, Kassen R. The fitness costs of antibiotic resistance mutations. Evol Appl. 2015;8(3):273–283. doi:10.1111/eva.12196.

9. Opatowski L, Varon E, Dupont C, Temime L, van der Werf S, Gutmann L, et al. Assessing pneumococcal meningitis association with viral respiratory infections and antibiotics: insights from statistical and mathematical models. Proceedings Biological Sciences. 2013;280(1764):20130519. doi:10.1098/rspb.2013.0519.

10. Boucher HW, Corey GR. Epidemiology of Methicillin-Resistant Staphylococcus aureus. Clin Infect Dis. 2008;46(Supplement 5):S344–S349. doi:10.1086/533590.

11. Cobey S, Lipsitch M. Niche and neutral effects of acquired immunity permit coexistence of pneumococcal serotypes. Science. 2012;335(6074):1376–1380. doi:10.1126/science.1215947.

12. Murall CL, Bauch CT, Day T. Could the human papillomavirus vaccines drive virulence evolution? Proc R Soc B. 2015;282(1798):20141069. doi:10.1098/rspb.2014.1069.

13. Aanensen DM, Feil EJ, Holden MTG, Dordel J, Yeats CA, Fedosejev A, et al. Whole-Genome Sequencing for Routine Pathogen Surveillance in Public Health: a Population Snapshot of Invasive Staphylococcus aureus in Europe. mBio. 2016;7(3):e00444–16. doi:10.1128/mBio.00444-16.

14. Lemey P, Rambaut A, Bedford T, Faria N, Bielejec F, Baele G, et al. Unifying Viral Genetics and Human Transportation Data to Predict the Global Transmission Dynamics of Human Influenza H3N2. PLoS Pathog. 2014;10(2):e1003932. doi:10.1371/journal.ppat.1003932.

15. Donker T, Reuter S, Scriberras J, Reynolds R, Brown NM, Török ME, et al. Population genetic structuring of methicillin-resistant Staphylococcus aureus clone EMRSA-15 within UK reflects patient referral patterns. Microbial Genomics. 2017;3(7). doi:10.1099/mgen.0.000113.

16. Obadia T, Silhol R, Opatowski L, Temime L, Legrand J, Thiébaut ACM, et al. Detailed Contact Data and the Dissemination of Staphylococcus aureus in Hospitals. PLOS Computational Biology. 2015;11(3):e1004170. doi:10.1371/journal.pcbi.1004170.

17. Nekkab N, Astagneau P, Temime L, Crépey P. Spread of hospital-acquired infections: A comparison of healthcare networks. PLOS Computational Biology. 2017;13(8):e1005666. doi:10.1371/journal.pcbi.1005666.

18. Fraser C, Hanage WP, Spratt BG. Neutral microepidemic evolution of bacterial pathogens. Proceedings of the National Academy of Sciences of the United States of America. 2005;102(6):1968–1973. doi:10.1073/pnas.0406993102.

19. Robinson K, Fyson N, Cohen T, Fraser C, Colijn C. How the Dynamics and Structure of Sexual Contact Networks Shape Pathogen Phylogenies. PLOS Computational Biology. 2013;9(6):e1003105. doi:10.1371/journal.pcbi.1003105.

20. Matt J Keeling PR. Modeling Infectious Diseases in Humans and Animals. Princeton University Press; 2008.

21. Wesolowski A, Eagle N, Tatem AJ, Smith DL, Noor AM, Snow RW, et al. Quantifying the impact of human mobility on malaria. Science (New York, NY). 2012;338(6104):267–270. doi:10.1126/science.1223467.

22. Balcan D, Colizza V, Gonçalves B, Hu H, Ramasco JJ, Vespignani A. Multiscale mobility networks and the spatial spreading of infectious diseases. Proceedings of the National Academy of Sciences of the United States of America. 2009;106(51):21484–21489. doi:10.1073/pnas.0906910106.

23. Louail T, Lenormand M, Picornell M, Cantú OG, Herranz R, Frias-Martinez E, et al. Uncovering the spatial structure of mobility networks. Nature Communications. 2015;6:6007. doi:10.1038/ncomms7007.

24. Rocha LEC, Liljeros F, Holme P. Information dynamics shape the sexual networks of Internet-mediated prostitution. Proceedings of the National Academy of Sciences. 2010;107(13):5706–5711. doi:10.1073/pnas.0914080107.

25. Mastrandrea R, Fournet J, Barrat A. Contact Patterns in a High School: A Comparison between Data Collected Using Wearable Sensors, Contact Diaries and Friendship Surveys. PLOS ONE. 2015;10(9):e0136497. doi:10.1371/journal.pone.0136497.

26. Salathé M, Kazandjieva M, Lee JW, Levis P, Feldman MW, Jones JH. A high-resolution human contact network for infectious disease transmission. Proceedings of the National Academy of Sciences. 2010;107(51):22020–22025. doi:10.1073/pnas.1009094108.

27. Génois M, Vestergaard CL, Fournet J, Panisson A, Bonmarin I, Barrat A. Data on face-to-face contacts in an office building suggest a low-cost vaccination strategy based on community linkers. Network Science. 2015;3(3):326–347. doi:10.1017/nws.2015.10.

28. Duval A, Obadia T, Martinet L, Boëlle PY, Fleury E, Guillemot D, et al. Measuring dynamic social contacts in a rehabilitation hospital: effect of wards, patient and staff characteristics. Scientific Reports. 2018;8(1):1686. doi:10.1038/s41598-018-20008-w.

29. Vanhems P, Barrat A, Cattuto C, Pinton JF, Khanafer N, Régis C, et al. Estimating Potential Infection Transmission Routes in Hospital Wards Using Wearable Proximity Sensors. PLOS ONE. 2013;8(9):e73970. doi:10.1371/journal.pone.0073970.

30. Voirin N, Payet C, Barrat A, Cattuto C, Khanafer N, Régis C, et al. Combining High-Resolution Contact Data with Virological Data to Investigate Influenza Transmission in a Tertiary Care Hospital. Infection Control & Hospital Epidemiology. 2015;36(3):254–260. doi:10.1017/ice.2014.53.

31. Assab R, Nekkab N, Crépey P, Astagneau P, Guillemot D, Opatowski L, et al. Mathematical models of infection transmission in healthcare settings: recent advances from the use of network structured data. Current Opinion in Infectious Diseases. 2017;30(4):410–418. doi:10.1097/QCO.0000000000000390.

32. Holme P, Saramäki J, editors. Temporal Networks. Springer; 2013.

33. Pastor-Satorras R, Castellano C, Van Mieghem P, Vespignani A. Epidemic processes in complex networks. Reviews of Modern Physics. 2015;87(3):925–979. doi:10.1103/RevModPhys.87.925.

34. Perra N, Gonçalves B, Pastor-Satorras R, Vespignani A. Activity driven modeling of time varying networks. Scientific Reports. 2012;2:469. doi:10.1038/srep00469.

35. Barthélemy M, Barrat A, Pastor-Satorras R, Vespignani A. Dynamical patterns of epidemic outbreaks in complex heterogeneous networks. Journal of Theoretical Biology. 2005;235(2):275–288. doi:10.1016/j.jtbi.2005.01.011.

36. Salathé M, Jones JH. Dynamics and Control of Diseases in Networks with Community Structure. PLOS Computational Biology. 2010;6(4):e1000736. doi:10.1371/journal.pcbi.1000736.

37. Sah P, Leu ST, Cross PC, Hudson PJ, Bansal S. Unraveling the disease consequences and mechanisms of modular structure in animal social networks. Proceedings of the National Academy of Sciences. 2017;114(16):4165–4170. doi:10.1073/pnas.1613616114.

38. Holme P, Liljeros F. Birth and death of links control disease spreading in empirical contact networks. Scientific Reports. 2014;4:4999. doi:10.1038/srep04999.

39. Darabi Sahneh F, Scoglio C. Competitive epidemic spreading over arbitrary multilayer networks. Physical Review E. 2014;89(6):062817. doi:10.1103/PhysRevE.89.062817.

40. Poletto C, Meloni S, Van Metre A, Colizza V, Moreno Y, Vespignani A. Characterising two-pathogen competition in spatially structured environments. Scientific Reports. 2015;5:7895. doi:10.1038/srep07895.

41. Poletto C, Meloni S, Colizza V, Moreno Y, Vespignani A. Host Mobility Drives Pathogen Competition in Spatially Structured Populations. PLOS Computational Biology. 2013;9(8):e1003169. doi:10.1371/journal.pcbi.1003169.

42. Sanz J, Xia CY, Meloni S, Moreno Y. Dynamics of Interacting Diseases. Phys Rev X. 2014;4(4):041005. doi:10.1103/PhysRevX.4.041005.

43. Karrer B, Newman MEJ. Competing epidemics on complex networks. Phys Rev E. 2011;84(3):036106. doi:10.1103/PhysRevE.84.036106.

44. Cai W, Chen L, Ghanbarnejad F, Grassberger P. Avalanche outbreaks emerging in cooperative contagions. Nature Physics. 2015;11(11):936–940. doi:10.1038/nphys3457.

45. Hébert-Dufresne L, Althouse BM. Complex dynamics of synergistic coinfections on realistically clustered networks. Proceedings of the National Academy of Sciences. 2015;112(33):10551–10556. doi:10.1073/pnas.1507820112.

46. Leventhal GE, Hill AL, Nowak MA, Bonhoeffer S. Evolution and emergence of infectious diseases in theoretical and real-world networks. Nature Communications. 2015;6:6101. doi:10.1038/ncomms7101.

47. Eames KTD, Keeling MJ. Coexistence and Specialization of Pathogen Strains on Contact Networks. The American Naturalist. 2006;168(2):230–241. doi:10.1086/505760.

48. Ballegooijen WMv, Boerlijst MC. Emergent trade-offs and selection for outbreak frequency in spatial epidemics. Proceedings of the National Academy of Sciences. 2004;101(52):18246–18250. doi:10.1073/pnas.0405682101.

49. Lion S, Gandon S. Spatial evolutionary epidemiology of spreading epidemics. Proceedings Biological Sciences. 2016;283(1841). doi:10.1098/rspb.2016.1170.

50. Buckee CO, Koelle K, Mustard MJ, Gupta S. The effects of host contact network structure on pathogen diversity and strain structure. PNAS. 2004;101(29):10839–10844. doi:10.1073/pnas.0402000101.

51. Buckee C, Danon L, Gupta S. Host community structure and the maintenance of pathogen diversity. Proceedings of the Royal Society of London B: Biological Sciences. 2007;274(1619):1715–1721. doi:10.1098/rspb.2007.0415.

52. Morris EK, Caruso T, Buscot F, Fischer M, Hancock C, Maier TS, et al. Choosing and using diversity indices: insights for ecological applications from the German Biodiversity Exploratories. Ecol Evol. 2014;4(18):3514–3524. doi:10.1002/ece3.1155.

53. Azaele S, Suweis S, Grilli J, Volkov I, Banavar JR, Maritan A. Statistical mechanics of ecological systems: Neutral theory and beyond. Rev Mod Phys. 2016;88(3):035003. doi:10.1103/RevModPhys.88.035003.

54. Kogan O, Khasin M, Meerson B, Schneider D, Myers CR. Two-strain competition in quasineutral stochastic disease dynamics. Phys Rev E. 2014;90(4):042149. doi:10.1103/PhysRevE.90.042149.

55. Ganesh A, Massoulie L, Towsley D. The effect of network topology on the spread of epidemics. In: Proceedings IEEE 24th Annual Joint Conference of the IEEE Computer and Communications Societies.. vol. 2; 2005. p. 1455–1466 vol. 2.

56. DeBenedictis PA. On the Correlations between Certain Diversity Indices. The American Naturalist. 1973;107(954):295–302. doi:10.1086/282831.

57. Pastor-Satorras R, Castellano C. Distinct types of eigenvector localization in networks. Scientific Reports. 2016;6:18847. doi:10.1038/srep18847.

58. Suweis S, Bertuzzo E, Mari L, Rodriguez-Iturbe I, Maritan A, Rinaldo A. On species persistence-time distributions. Journal of Theoretical Biology. 2012;303:15–24. doi:10.1016/j.jtbi.2012.02.022.

59. van Kleef E, Robotham JV, Jit M, Deeny SR, Edmunds WJ. Modelling the transmission of healthcare associated infections: a systematic review. BMC Infect Dis. 2013;13:294. doi:10.1186/1471-2334-13-294.

60. Martinet L, Crespelle C, Fleury E, Boëlle PY, Guillemot D. The Link Stream of Contacts in a Whole Hospital. arXiv:180505752 [cs]. 2018;.

61. Chang Q, Lipsitch M, Hanage WP. Impact of Host Heterogeneity on the Efficacy of Interventions to Reduce Staphylococcus aureus Carriage. Infect Control Hosp Epidemiol. 2016;37(2):197–204. doi:10.1017/ice.2015.269.

62. Kouyos R, Klein E, Grenfell B. Hospital-Community Interactions Foster Coexistence between Methicillin-Resistant Strains of Staphylococcus aureus. PLOS Pathogens. 2013;9(2):e1003134. doi:10.1371/journal.ppat.1003134.

63. Bonten MJ, Austin DJ, Lipsitch M. Understanding the spread of antibiotic resistant pathogens in hospitals: mathematical models as tools for control. Clinical Infectious Diseases: An Official Publication of the Infectious Diseases Society of America. 2001;33(10):1739–1746. doi:10.1086/323761.

64. Margolis E, Yates A, Levin BR. The ecology of nasal colonization of Streptococcus pneumoniae, Haemophilus influenzae and Staphylococcus aureus: the role of competition and interactions with host’s immune response. BMC Microbiol. 2010;10:59. doi:10.1186/1471-2180-10-59.

65. Dall’Antonia M, Coen PG, Wilks M, Whiley A, Millar M. Competition between methicillin-sensitive and - resistant Staphylococcus aureus in the anterior nares. J Hosp Infect. 2005;61(1):62–67. doi:10.1016/j.jhin.2005.01.008.

66. Cobey S, Baskerville EB, Colijn C, Hanage W, Fraser C, Lipsitch M. Host population structure and treatment frequency maintain balancing selection on drug resistance. Journal of The Royal Society Interface. 2017;14(133):20170295. doi:10.1098/rsif.2017.0295.

67. van Duijkeren E, Ikawaty R, Broekhuizen-Stins MJ, Jansen MD, Spalburg EC, de Neeling AJ, et al. Transmission of methicillin-resistant Staphylococcus aureus strains between different kinds of pig farms. Veterinary Microbiology. 2008;126(4):383–389. doi:10.1016/j.vetmic.2007.07.021.

68. Gjini E, Valente C, Sá-Leão R, Gomes MGM. How direct competition shapes coexistence and vaccine effects in multi-strain pathogen systems. Journal of Theoretical Biology. 2016;388:50–60. doi:10.1016/j.jtbi.2015.09.031.

69. Lipsitch M, Colijn C, Cohen T, Hanage WP, Fraser C. No coexistence for free: Neutral null models for multistrain pathogens. Epidemics. 2009;1(1):2–13. doi:10.1016/j.epidem.2008.07.001.

70. Buendía V, Muñoz MA, Manrubia S. Limited role of spatial self-structuring in emergent trade-offs during pathogen evolution. arXiv:180408463 [q-bio]. 2018;.

71. Sofonea MT, Alizon S, Michalakis Y. Exposing the diversity of multiple infection patterns. Journal of Theoretical Biology. 2017;419:278–289. doi:10.1016/j.jtbi.2017.02.011.

72. Li M, Rao VD, Gernat T, Dankowicz H. Lifetime-preserving reference models for characterizing spreading dynamics on temporal networks. Scientific Reports. 2018;8(1):709. doi:10.1038/s41598-017-18450-3.

73. Stanoev A, Trpevski D, Kocarev L. Modeling the Spread of Multiple Concurrent Contagions on Networks. PLOS ONE. 2014;9(6):e95669. doi:10.1371/journal.pone.0095669.

74. Lauderdale TLY, Wang JT, Lee WS, Huang JH, McDonald LC, Huang IW, et al. Carriage rates of methicillin-resistant *Staphylococcus aureus* (MRSA) depend on anatomic location, the number of sites cultured, culture methods, and the distribution of clonotypes. Eur J Clin Microbiol Infect Dis. 2010;29(12):1553–1559. doi:10.1007/s10096-010-1042-8.

75. Gardiner C. Stochastic Methods: A Handbook for the Natural and Social Sciences. 4th ed. Springer Series in Synergetics. Berlin Heidelberg: Springer-Verlag; 2009.

